# Subconfluent ARPE-19 Cells Display Mesenchymal Cell-State Characteristics and Behave Like Fibroblasts, Rather than Epithelial Cells, in Experimental HCMV Infection Studies

**DOI:** 10.1101/2023.10.27.564463

**Authors:** Preethi Golconda, Mariana Andrade-Medina, Adam Oberstein

## Abstract

Human cytomegalovirus (HCMV) has a broad cellular tropism and epithelial cells are important physiological targets during infection. The retinal pigment epithelial cell line ARPE-19 has been used to model HCMV infection in epithelial cells for decades and remains a commonly used cell-type for studying viral entry, replication, and the cellular response to infection. We previously found that ARPE-19 cells, despite being derived from an epithelial cell explant, express extremely low levels of canonical epithelial proteins, such as E-cadherin and EpCAM. Here, we perform comparative studies of ARPE-19 and additional epithelial cell lines with strong epithelial characteristics. We find that ARPE-19 cells cultured under subconfluent conditions resemble mesenchymal fibroblasts, rather than epithelial cells; consistent with previous studies showing that ARPE-19 cultures require extended periods of high confluency culture to maintain epithelial characteristics. By reanalyzing public gene expression data and using machine-learning, we find evidence that ARPE-19 cultures maintained across many labs exhibit mesenchymal characteristics and that the majority of studies employing ARPE-19 use them in a mesenchymal state. Lastly, by performing experimental HCMV infections across mesenchymal and epithelial cell lines, we find that ARPE-19 cells behave like mesenchymal fibroblasts, producing logarithmic yields of cell-free infectious progeny, while cell lines with strong epithelial character exhibit an atypical infectious cycle and naturally restrict the production of cell-free progeny. Our work highlights important characteristics of the ARPE-19 cell line and suggests that sub-confluent ARPE-19 cells may not be optimal for modeling epithelial infection with HCMV or other human viruses. It also suggests that HCMV biosynthesis and/or spread may occur quite differently in epithelial cells compared to mesenchymal cells. These differences could contribute to viral persistence or pathogenesis in epithelial tissues.

## Introduction

Human cytomegalovirus (HCMV), a betaherpesvirus, is an important human pathogen with incompletely understood biology. The virus causes birth defects in newborns, morbidity in transplant patients, and is increasingly linked to chronic diseases, such as cancer or immune senescence. A vaccine for HCMV is not available and greater knowledge about HCMV biology is needed for continued therapeutic development.

Epithelial cells serve as primary targets for HCMV in the human body, playing pivotal roles in viral entry, dissemination, and the development of CMV-induced inflammatory diseases in various organs [1,2], including the intestine (CMV gastroenteritis), lungs (CMV pneumonia), and eyes (CMV retinitis) [1–4]. Furthermore, HCMV is suspected to establish persistent, possibly immunologically silent, infections in epithelial cells of the breast, kidney, and oral cavity [2,5–8].

The cell line ARPE-19 [9], derived from a primary human adult retinal pigment epithelium (RPE) explant, is commonly used to study HCMV in epithelial cells. ARPE-19 cells are not immortalized [9], but can be expanded substantially, thus providing a fairly stable source of retinal epithelial cells for functional and genetic studies. Additionally, they are both susceptible and permissive to HCMV infection [10,11] which makes them useful for studying mechanisms of entry, replication [12–17], and cellular responses to infection [10,18]. Since the virus is found in RPE cells during CMV retinitis [2,3,19–21], experimentally infected ARPE-19 cells are seen as a physiologically relevant *in-vitro* model for studying HCMV in epithelial cells [22].

The majority of recent studies using ARPE-19 for modeling HCMV infection (including our own [10,11]) adsorb the virus onto just confluent or slightly subconfluent (e.g., 95% confluent) monolayers. However, ARPE-19 cells display density-dependent changes in their biochemistry [9,23,24]. For example, gene expression studies have shown large increases in the expression of visual cycle, melanogenesis, and epithelial junction genes after extended (four months), high-confluency culture [23]. Extended, high confluency culture is also required for formation of tight-junctions, establishment of apico-basal polarization, and development of barrier function [9,25,25,26]. This functionally differentiated, high density ARPE-19 state most resembles primary RPE [23,27]. Thus, it is possible that the low confluency ARPE-19 cell state commonly used to study HCMV differs significantly from the physiologic epithelial state the virus encounters during natural infection in the eye.

In a previous study, we conducted comparative experimental infections with HCMV across ARPE-19 cells and permissive MRC-5 fibroblasts [10]. To our surprise, we observed that E-cadherin [28–31] and EpCAM [32–34], two canonical epithelial markers, were undetectable in uninfected ARPE-19 cells (see Figure 6E in [10]). Additionally, comprehensive transcriptomics analysis was unable to identify any significant cell-type specific responses to HCMV infection. These observations suggested that ARPE-19 cells were phenotypically similar to fibroblasts, rather than epithelial cells, raising concerns about their reliability as an epithelial infection model for HCMV. Given the known effects of cell-density on ARPE-19 biology, we hypothesized that low confluency culture (subculturing at 95 % confluency) might encourage these cells to adopt a mesenchymal, rather than an epithelial, cell state.

Employing computational and genetic approaches, we have found that ARPE-19 cells maintained at sub-confluency do indeed exhibit a mesenchymal, fibroblast-like phenotype that distinguishes them from typical epithelial cells. These properties are not specific to ARPE-19 cultures maintained in our laboratory, since re-analysis of RNA-sequencing data from a number of different laboratories show nearly identical results. Genetic experiments involving epithelial and mesenchymal transcription factors have revealed that ARPE-19 cells possess limited plasticity to transition toward a further mesenchymal cell state but can readily transition to an epithelial state. This finding supports the notion that their baseline phenotype under subconfluent culture conditions tilts toward the mesenchymal end of the epithelial-mesenchymal axis. Furthermore, consistent with observations in the literature [23], prolonged growth at high confluency stimulated epithelial and RPE-specific gene expression, highlighting the dependency of epithelial features on cell density. However, long-term culture led to increased gene expression of both epithelial and mesenchymal genes, suggesting that even at high confluency, ARPE-19 cells may maintain a hybrid, potentially aberrant epithelial-mesenchymal cell state. Finally, by performing comparative experimental infections across ARPE-19 and strongly epithelial cells lines, we have found that ARPE-19 cells produce high yields of infectious progeny, similar to fibroblasts, while epithelial cell lines with strong epithelial characteristics display an atypical infectious cycle where cell-free infectious progeny production is strongly restricted.

Our findings underscore the significance of the epithelial-mesenchymal cell state axis as a potential modulator of HCMV infection. They also suggest that caution be exercised when using ARPE-19 cells to model epithelial infection with HCMV or other human viruses. We propose that the epithelial-mesenchymal cell state axis may serve as an intrinsic regulator of HCMV infection in epithelial cells, potentially influencing viral biosynthesis, spread, or immune evasion; each of which could impact viral persistence or pathogenesis in epithelial tissues.

## Materials and Methods

### Cell Lines, Culture Conditions, and Viruses

MRC-5 embryonic lung fibroblasts, ARPE-19 adult retinal pigment epithelial cells, MCF10A mammary epithelial cells, RWPE-1 prostate epithelial cells were obtained from the American Type Culture Collection. MRC-5 and ARPE-19 cells were grown in DMEM supplemented with 10% fetal bovine serum, 1 mM sodium pyruvate, 2 mM glutamax (Gibco), 10 mM Hepes pH 7.4, 0.1 mM MEM Non-Essential Amino Acids (Gibco), 100 units/ml Penicillin G, and 100 μg/ml Streptomycin Sulfate. MCF10A cells were cultured in DMEM supplemented with 5% horse serum, 10 mM Hepes pH 7.4, 20 ng/ml EGF, 10 ug/ml Insulin, 1 nM Forskolin, 500 ng/ml Hydrocortisone, 100 units/ml Penicillin G, and 100 μg/ml Streptomycin Sulfate. RWPE-1 cells were cultured in K-SFM (Gibco) supplemented with 25 units/ml Penicillin G and 25 μg/ml Streptomycin Sulfate. HCMV strain TB40/E-BAC4 [35] was reconstituted by electroporation into ARPE-19 cells. Virus stocks were titered on MRC-5 and ARPE-19 cells by an HCMV infectious unit assay (see below).

### RNA-sequencing Data Re-analysis

Public gene expression data was collected from the Gene Expression Omnibus (GEO) repository [36] (https://www.ncbi.nlm.nih.gov/geo/). Nine to eleven independent studies containing untreated or control treated samples of MCF10A, RWPE-1, ARPE-19, and MRC-5 cells were manually identified and metadata from each study (Table S1) was uniformly curated to allow for automated download and processing. Raw reads from RNA-sequencing fastq files were pseudo-aligned to the *homo sapiens* transcriptome and converted to pseudo-counts using Kallisto [37]. Human genome release 34 (GRCh38.p13) was used for this analysis and reference transcripts and annotation metadata (gff3) were acquired from Genecode (https://www.gencodegenes.org/human/release_34.html). The count matrices for all runs were joined and normalized for transcript length (within sample normalization) and compositional bias (cross-sample normalization) using the GeTMM [38] procedure and the *R*-package limma [39,40].

Hierarchical clustering was performed using the R-functions *hclust* from the R-package gplots [41] with method=“ward.D2” and distfun=function(x) as.dist(1-cor(t(x), method=“spearman”)). Heatmaps were generated using the function *heatmap*.*2* from the *R*-package gplots [41] and ggplot2 [42]. Cell-type classification (epithelial vs mesenchymal prediction) was performed in *R*[43] using the MLseq [44] and caret [45] packages. Six different binary classifiers (Table 1) were trained on the GeTMM normalized count data for MCF10A, RWPE-1, and MRC-5, excluding ARPE-19 data; with MCF10A and RWPE-1 being assigned an “epithelial” (E) class label and MRC-5 being assigned a “mesenchymal” (M) class label. Tuning parameters were optimized using 5-fold cross-validation repeated 10 times. Accuracy was assessed by training on 70% of the input data (still excluding ARPE-19) and predicting the classes of the remaining 30%. All classifiers achieved 100%% accuracy. Lastly, each trained model was used to predict the class of each ARPE-19 RNA-sequencing data set, using the top 5000 most variable genes.

**Table 1.** Prediction of the ARPE-19 Phenotype Using Machine-Learning Classification. Public gene expression data from Figures 1 and 2 were used to train six binary classifiers as described in “Materials and Methods.” The proportion of RNA-seq runs predicted to be epithelial (“E”) or mesenchymal (“M”) is listed at the bottom of the table. Run IDs with hyphenated accession numbers are technical replicates used to increase RNA-sequencing coverage, which were concatenated and considered a single pooled sample.

| Predicted Phenotypic Class (E = Epithelial; M = Mesenchymal) |  |  |  |  |  |  |  |
| --- | --- | --- | --- | --- | --- | --- | --- |
| RNA-seq Run ID | Cell Type | KNN | Logistic Regression | Naive Bayes | Support Vector Machine (Linear) | Support Vector Machine (Polynomial) | Random Forrest |
| SRR5807527, -28, -29, -30 | ARPE-19 | M | M | M | M | M | M |
| SRR5807531, -32, -33, -34 | ARPE-19 | M | M | M | M | M | M |
| SRR5807535, -36, -37, -38 | ARPE-19 | M | M | M | M | M | M |
| SRR12193639 | ARPE-19 | M | M | M | M | M | M |
| SRR12193640 | ARPE-19 | M | M | M | M | M | M |
| SRR12193641 | ARPE-19 | M | M | M | M | M | M |
| SRR19895498, -99 | ARPE-19 | E | M | E | M | M | E |
| SRR19895500, -01 | ARPE-19 | M | M | M | M | M | M |
| SRR19895502, -03 | ARPE-19 | M | M | M | M | M | M |
| SRR5629569, -70 | ARPE-19 | M | M | M | M | M | M |
| SRR5629579, -80 | ARPE-19 | M | M | M | M | M | M |
| SRR19033104 | ARPE-19 | M | M | M | M | M | M |
| SRR19033105 | ARPE-19 | M | M | M | M | M | M |
| SRR19033106 | ARPE-19 | M | M | M | M | M | M |
| SRR8032314 | ARPE-19 | M | M | M | M | M | M |
| SRR8032315 | ARPE-19 | M | M | M | M | M | M |
| SRR8032316, -17 | ARPE-19 | M | M | M | M | M | M |
| SRR10766046 | ARPE-19 | E | M | M | M | M | M |
| SRR10766047 | ARPE-19 | E | M | M | M | M | M |
| SRR10766048 | ARPE-19 | E | M | M | M | M | M |
| SRR4427911 | ARPE-19 | M | M | M | M | M | M |
| SRR4427912 | ARPE-19 | M | M | M | M | M | M |
| SRR4427913 | ARPE-19 | M | M | M | M | M | M |
| SRR12795670 | ARPE-19 | M | M | M | M | M | M |
| SRR12795671 | ARPE-19 | M | M | M | M | M | M |
| Proportion Mesenchymal: |  | 0.84 | 1.00 | 0.96 | 1.00 | 1.00 | 0.96 |

### HCMV Infectious Unit Assay

Supernatants from infected cells were collected, centrifuged at 500 x*g* for 5 min to remove cellular debris, and serially diluted into complete media. For stock titration, 50 ul of concentrated HCMV particles were 10-fold serially diluted into virus storage buffer (PBS + 7% sucrose + 1% BSA) and 100 ul of each dilution was used to infect confluent monolayers of MRC-5 fibroblasts and ARPE-19 cells seeded into 24-well plates. At 20 hpi, cells were fixed with 100% methanol, stained for HCMV immediate early-1 (1E1) antigen using monoclonal antibody 1B12 [46], and imaged using a Keyence BZ-X710 Inverted Microscope. Wells with 20% or less infected nuclei were selected to avoid saturation effects. IE1-positive nuclei in each image were counted using CellProfiler [47] v2.1.1. The mean titer and standard deviation for each condition was calculated by multiplying the number of IE1-positive nuclei on each image by (1/dilution factor), (1.0 ml/infection volume) in milliliters, and a well factor. The well-factor was the total well surface area (acquired from the plate manufacturer’s engineering specification) divided by the image area (calculated using a calibration slide and the image [48] function “set scale”). Four images/well (technical replicates) were sampled from three independent infections (biological replicates). Infectious units per mL (IU/mL) were determined by first averaging the three technical replicates for each biological replicate (using the equation described above), converting each average to a titer, and then calculating the average and standard deviation of the biological replicates.

### HCMV Growth Curves

Two days prior to infection, RWPE-1 cells were seeded into 24-well plates at 3e5/well. One day prior to infection MCF10A, ARPE-19, and MRC-5 cells were seeded into 24-well plates at 2e5/well, 2e5/well, and 2.5e5/well, respectively. One 24-well plate per time point was seeded with triplicate wells of each cell type. The day of infection, cells were counted and media was changed to 0.7 ml complete media specific for each cell type (see “Materials and Methods: Cell Lines, Culture Conditions, and Viruses”). An identical HCMV virus stock was then added to triplicate wells of each cell type at an MOI of 1 IU/cell (MRC-5, ARPE-19, RWPE-1) or 3 IU/cell (MCF10A). Cultures were incubated at 37°C/5% CO_2_ for 2 hours with gentle tapping every 30 minutes. At 2 hpi, monolayers were washed one time with warm PBS, 0.8 ml complete medium was added to each well, and cells were returned to the tissue culture incubator. At 1, 2, 3, 4, 5, and 6 days post-infection (dpi), cell supernatants were collected, cleared of detached cells by centrifuging at 300 x*g* for 5 min at RT, aliquoted, and frozen at -80°C until all time points were collected. Infected cell monolayers were washed with ice-cold PBS and harvested for western analysis (see “Materials and Methods: “Western Blotting”). Once all time points were harvested, supernatants were thawed (1x freeze-thaw cycle), serially diluted in DMEM/10% FBS in 96-well plates, and adsorbed onto confluent monolayers of MRC-5 fibroblasts. 24 hours later, reporter fibroblasts were fixed with cold MeOH and cell-free infectious progeny titers were determined using an infectious unit assay (see “Materials and Methods: HCMV Infectious Unit Assay”).

### HCMV Entry Assay

The percentage of IE1-positive cells at 24 hpi was used to assess the frequency of infection with HCMV across the different cell lines, at different multiplicities of infection. Cells were seeded into 24-well plates in biological triplicate for each MOI and infected with HCMV as described above (see “Materials and Methods: “HCMV Growth Curves”). At 24 hpi, monolayers were fixed in cold MeOH and stained for IE1 antigen using monoclonal antibody 1B12 [46]. Cells were counterstained with DAPI to visualize nuclei and the percentage of infected cells in each condition was calculated using fluorescence imaging. For quantification, nine random fields (technical replicates) were imaged in the IE1 and DAPI channels, in each well. The percentage of infected cells in each field was calculated by dividing the number of IE1-positive nuclei by the total number of DAPI-positive, which were counted using CellProfiler [47] v2.1.1 software. The nine technical replicates were averaged to yield an estimate of the mean percentage of infected cells in each well, and then the “mean-of-means” and standard deviation were calculated across three independent infections (biological replicates) for each condition.

### Vector and Stable Cell Line Construction

Sequence confirmed cDNAs encoding Snail and OVOL2 were obtained from the DNASU plasmid repository (Arizona State University) and subcloned into pLVX-TetOne-Puro (Takara) vector. Lentiviral vectors were produced by co-transfecting each pLVX transfer vector with the packaging plasmids pCMV-dR8.91 and pCMV-VSV-G into HEK293FT cells. Plasmids were mixed in a 12:12:1 (vector:dR8.91:VSV-G) ratio by mass and mixed with an empirically optimized amount of branched polyethyleneimine (Sigma) before adding to cells. Lentivirus-containing supernatants were collected at 48 and 72 hours post transfection and concentrated over a 20% sorbitol cushion using ultracentrifugation. Lentivirus pellets were resuspended in PBS containing 7% Sucrose and 1% BSA. Transductions were performed by empirically determining the volume of lentivirus allowing 60 to 70 % of the cells to survive puromycin selection; an effective MOI of approximately 1.0. 5 ug/ml hexadimethrine bromide (“polybrene”; Sigma) was added during transduction. 48 hours after transduction, selective media containing 2 ug/ml puromycin was added until non-transduced cells were completely killed, at which point transduced cells were maintained in 1 ug/ml puromycin for expansion. To induce Snail and OVOL2, cell populations were treated with either 1 ug/ml doxycycline for two to four days.

### RNA-preparation and Quantitative Reverse-Transcription PCR (*q*RT-PCR) Analysis

For *q*RT-PCR analysis, cells were collected in Tri-reagent (Sigma) and total RNA was isolated using an RNeasy RNA isolation kit (Qiagen). DNA was removed using Turbo DNase (Thermo Fisher Scientific) and RNA was quantified using a NanoDrop Spectrophotometer (Thermo Scientific). cDNA was prepared using Superscript III reverse transcriptase (Invitrogen) and random hexamer primers. *q*RT-PCR reactions were performed using Power SYBR Green Master Mix (Applied biosystems) and data was collected on a Viia7 digital PCR machine (Life Technologies). Data was analyzed using the ΔΔCT method [49] with cyclophilin A (*PPIA*) as the reference gene. *CDH1* and *VIM* primers were from Mani et. al [50]. All other primers were designed using either QuantPrime [51] or GETPRime [52]. *q*RT-PCR primer sequences are listed in Supplementary Table 2.

### Western Blotting

For western analysis, cell monolayers were collected in RIPA buffer (50 mM Hepes pH 7.4, 150 mM NaCl, 1% NP-40, 0.5% sodium deoxycholate, 0.1% SDS, 5 ug/ml aprotinin, 10 ug/ml leupeptin, 1 mM PMSF). Samples were then sonicated and debris was cleared by centrifugation. Protein concentration was determined using the BCA assay (Pierce) and equal amounts of total protein were separated by SDS-PAGE. Proteins were transferred to PVDF membranes and blocked with 5% BSA in HBST (50 mM Hepes pH 7.4, 150 mM NaCl, 0.05% tween-20). Primary antibodies were diluted in 1% BSA and rocked at 4°C overnight. Primary antibodies were detected using HRP-conjugated secondary antibodies and ECL Prime (Cytiva). Antibodies and dilutions are listed in Supplementary Table 3.

### Data Availability Statement

Kallisto pseudo counts, study metadata, count matrices, and code used to perform normalization, clustering, machine-learning analyses, and data visualization are available at the following public github repository: https://github.com/aoberstein/2023-Golconda_A19_paper.git.

## Results

### The ARPE-19 Cell Line Exhibits Prominent Fibroblast-like, Mesenchymal Features

To corroborate the low baseline expression levels of E-cadherin and EpCAM in ARPE-19 cells and to gauge the expression range of these proteins in epithelial cells, we selected two widely employed epithelial cell lines from the biomedical literature as positive controls: MCF10A mammary epithelial cells [53] and RWPE-1 prostate epithelial cells [54]. MCF10A cells are spontaneously immortalized [53], while RWPE-1 cells have been immortalized using HPV-18 [54]. Both cell lines, however, remain untransformed and maintain stable epithelial morphology and functional characteristics *in-vitro* [53–60]. HCMV has been found in both mammary epithelial cells [7] and prostate epithelial cells [61,62] *in-vivo*; therefore, MCF10A and RWPE-1 are physiologically relevant experimental infection models. Our initial assessment involved comparing the steady-state expression levels of E-cadherin, EpCAM, and the mesenchymal marker Vimentin [63–66] across the epithelial control lines (MCF10A and RWPE-1), ARPE-19 cells, and MRC-5 fibroblasts - a primary lung embryonic fibroblast culture serving as a mesenchymal control. As anticipated from our previous observations, E-cadherin and EpCAM were undetectable in ARPE-19 cells and MRC-5 fibroblasts, while the epithelial controls expressed high levels of both epithelial markers (Fig. 1A). Conversely, steady-state levels of Vimentin, N-cadherin, and OB-cadherin were elevated in ARPE-19 and MRC-5 cells, yet undetectable in MCF10A and RWPE-1 (Fig. 1A). To further substantiate these findings at the mRNA level and assess the expression of a broader panel of epithelial and mesenchymal markers, we conducted gene expression analysis across these cell lines. Quantitative reverse-transcription PCR (*q*RT-PCR) analysis revealed the following: (1) exceptionally high expression levels of epithelial genes in MCF10A and RWPE-1 compared to MRC-5 and ARPE-19 cells (e.g., *CDH1*/*E-cadherin* mRNA was >4000-fold higher in MCF10A or RWPE-1 than in either MRC-5 or ARPE-19) (Fig. 1B); (2) significantly higher levels of mesenchymal genes (e.g. *VIM, FBN1, CDH11*) in ARPE-19 and MRC-5 relative to MCF10A and RWPE-1 (Fig. 1B); and (3) a gene expression pattern in ARPE-19 cells highly reminiscent of that in MRC-5 fibroblasts across most analyzed genes (Fig. 1B). Hence, our western blot (Fig. 1A) and *q*RT-PCR (Fig. 1B) analyses confirm that our ARPE-19 cells, analyzed immediately upon reaching confluency, assume a mesenchymal phenotype as opposed to an epithelial one.

**Figure 1.**
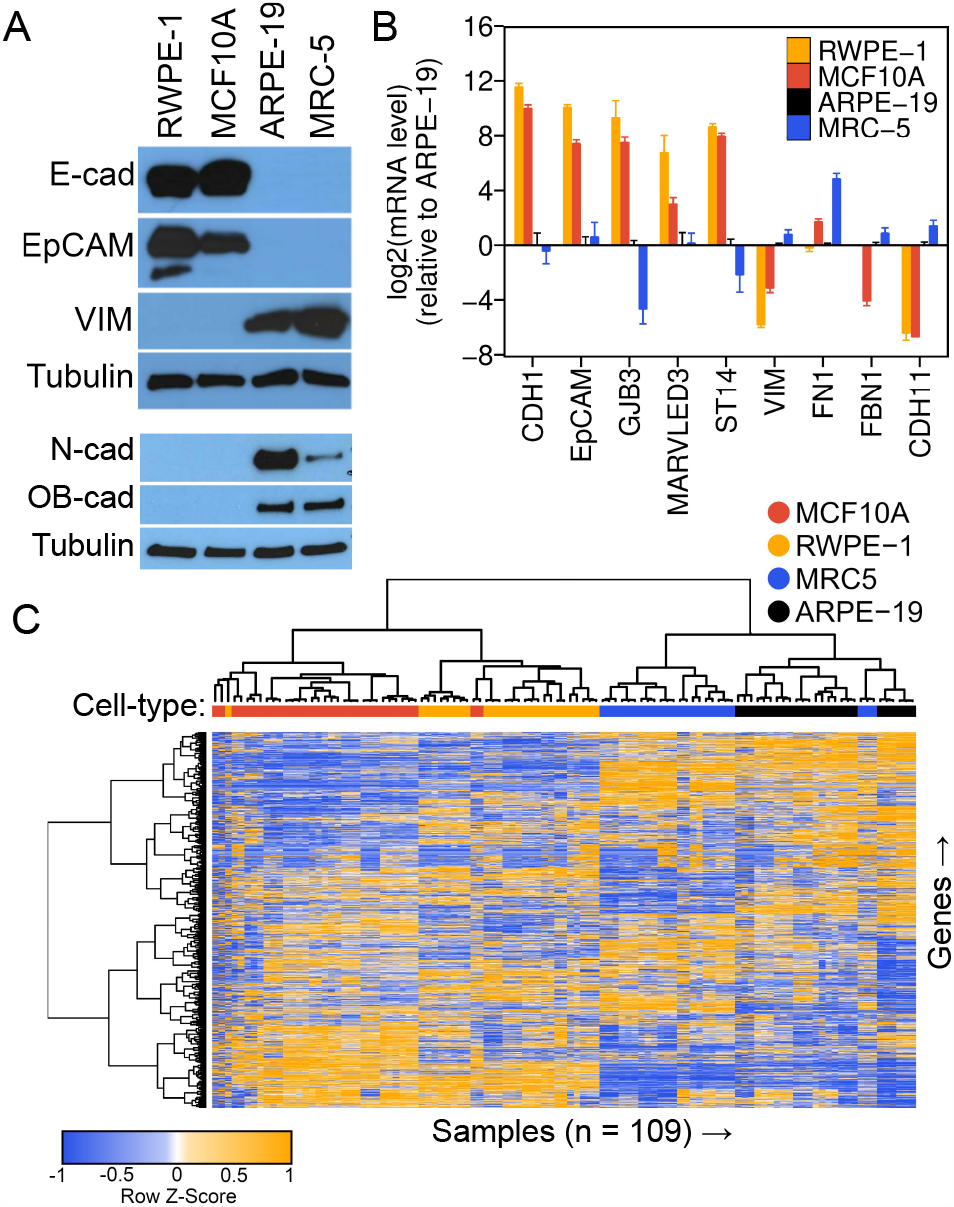
ARPE-19 Cells Display a Fibroblast-like, Mesenchymal Phenotype. (A) Western analysis of epithelial (E-cadherin, EpCAM) and mesenchymal (VIM, N-cadherin, OB-cadherin) marker proteins across RWPE-1, MCF10A, ARPE-19, and MRC-5 cell lines. ▫-Tubulin is shown as a loading control. (B) Relative mRNA levels of epithelial (*CDH1, EpCAM, GJB3, MARVELD3, ST14)* and mesenchymal (*VIM, FN1, FBN1, CDH11*) marker genes measured by *q*RT-PCR. Data represent the mean +/-SD from three biological replicates assayed in technical triplicate. (C) Unsupervised clustering analysis of public gene expression data. 109 samples from 38 RNA-sequencing studies (see *Table S1*) of untreated or control-treated RWPE-1, MCF10A, ARPE-19, and MRC-5 cells were uniformly processed, normalized, clustered using 1-minus the Spearman’s correlation coefficient as a distance metric (see “Materials and Methods”) and displayed as a heatmap with dendrograms showing related clusters. Samples from the same cell types formed clusters and ARPE-19 cells established a clade with MRC5 cells, suggesting a similar global gene expression pattern.

Cell lines can exhibit phenotypic variations when maintained in different laboratories [67]. To address the potential for phenotypic drift in our specific ARPE-19 culture and comprehensively examine the hypothesis that commonly cultured ARPE-19 cells exhibit a mesenchymal phenotype, we conducted gene expression and classification analyses using publicly available transcriptomics data from MCF10A, RWPE-1, MRC-5, and ARPE-19 cells. Our goal was to establish a consensus transcriptomic signature for ARPE-19 cells by compiling data from numerous independent studies conducted in different laboratories. This entailed collecting data from nine to eleven distinct RNA-sequencing studies for each cell type, all sourced from the Gene Expression Omnibus [36] (GEO) (Table S1). In order to maximize the number of data sets from different labs, some control samples from perturbation experiments were included (e.g., siRNA control or vector controls); however, care was taken to acquire as many untreated RNA-seq runs for each cell type, wherever feasible. We uniformly processed and normalized the 109 individual samples collected from these studies (see “Materials and Methods”) and conducted a comparative analysis on the resulting gene expression profiles. Our analysis comprised two primary components: (1) unsupervised hierarchical clustering to group samples based on their global gene expression patterns; and (2) supervised machine learning (ML) analysis to predict the epithelial-like (E) or mesenchymal-like (M) phenotype of ARPE-19 cells, employing ML-classifiers trained on experimentally derived epithelial and mesenchymal gene expression profiles.

The hierarchical clustering definitively separated the four cell types into distinct clades (Fig. 1C), indicating that the variation in gene expression due to batch effects, or other confounding variables, was minimal compared to the variation due to cell type identity. Two major phenotypic clades emerged, with ARPE-19 cells clustering closely with MRC-5 cells, and MCF10A cells clustering alongside RWPE-1 cells (Fig. 1C). Analysis of individual epithelial and mesenchymal marker transcripts revealed that ARPE-19 displayed a gene expression pattern akin to that of MRC-5, with expression levels closely resembling those of MRC-5 cells and exhibiting an inverse correlation with those of MCF10A or RWPE-1 cells (Fig. 2). For instance, *CDH1* and *EPCAM* exhibited extremely low mean expression levels (< 1 rpkm) in both ARPE-19 and MRC-5 cells, while robust expression (between 32 and 64 rpkm) was observed in MCF10A and RWPE-1 cells. Conversely, mesenchymal markers, such as *CDH11, FBN1, VIM, ZEB1*, and *ZEB2*, displayed an opposite trend (Fig. 2).

**Figure 2.**
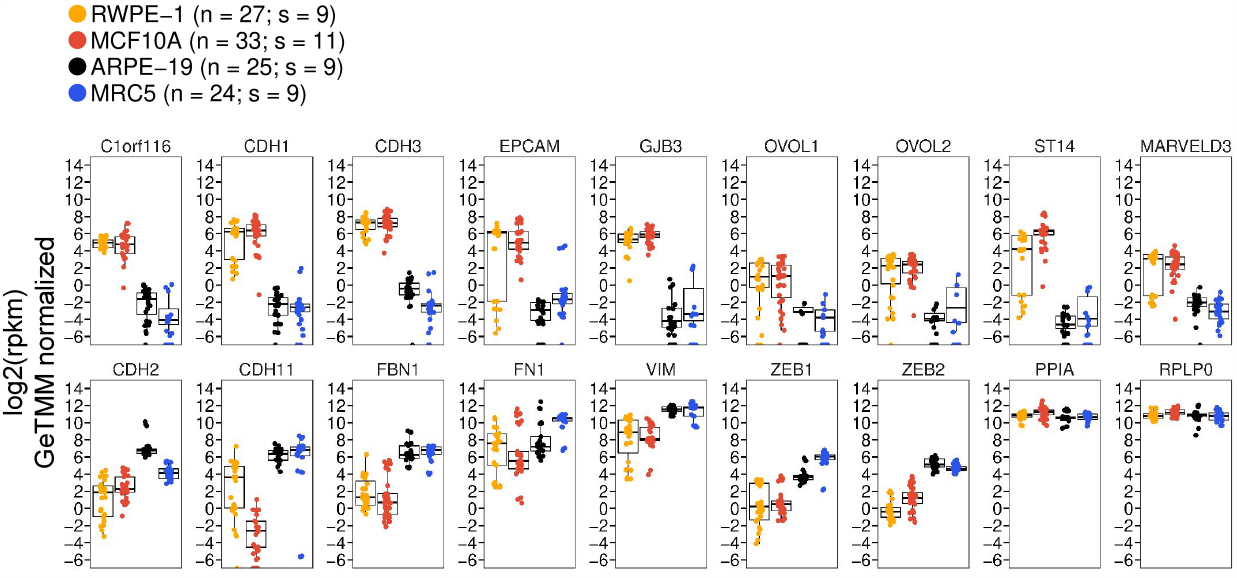
mRNA Expression Levels of Epithelial and Mesenchymal Genes from Public Transcriptomics Data. Relative expression levels of epithelial marker genes (upper panel), mesenchymal marker genes (lower panel, first seven plots), and normalization controls (*PPIA* and *RPLP0*). RNA-sequencing data was re-processed from the raw data from 38 studies, representing 109 sequencing runs (RWPE-19, *n* = 27; MCF10A, *n* = 33; ARPE-19, *n* = 25; MRC5, *n* = 24). Data represent the absolute expression level in reads-per-kilobase-per-million (rpkm) in each cell-type, after cross-sample normalization of the 109 runs using the GeTMM procedure (see “Material and Methods”). Data are shown as strip plots with overlayed box-plots showing the median and interquartile range in each cell type. *n* = number of sequencing runs; *s* = number of studies.

We next employed supervised ML techniques to predict the phenotypic class of ARPE-19 cells. For this analysis, we trained different binary ML classification models on a subset of the gene expression data used for clustering analysis (Table S1 and Fig. 1C). To train these models, we removed the ARPE-19 samples from the normalized gene expression matrix and assigned an “M” or “E” class to each remaining sample (MRC-5 = M; MCF10A and RWPE-1 = E). Algorithms were then trained on 70% of the remaining data, and predictive accuracy was assessed by predicting the E vs. M class of the remaining 30%, still excluding ARPE-19. All models achieved 100% accuracy on the training data, as determined using a resampling (cross-validation) procedure (see “Materials and Methods”). Each model was then used to predict the E vs M class of each ARPE-19 sample. Using six different commonly used binary classifiers, the vast majority of ARPE-19 samples were projected to be mesenchymal (Table 1). Three out of the six classifiers predicted all ARPE-19 samples to be mesenchymal, while the remaining three classifiers predicted 96%, 96%, or 84% of the samples to be mesenchymal. These results align with the hierarchical clustering analysis and further affirm that ARPE-19 cells exhibit a mesenchymal phenotype closely resembling that of MRC-5. In summary, our analysis of public gene expression data substantiates that ARPE-19 cells derived from multiple independent laboratories exhibit a mesenchymal phenotype akin to that of MRC-5, thereby validating our immunoblotting and *q*RT-PCR results characterizing our laboratory’s ARPE-19 cell stock.

### Sub-Confluent ARPE-19 Cells Are Inducible to an Epithelial Cell State but Not a Further Mesenchymal State

Most epithelial cells possess the capacity to transition between epithelial and mesenchymal cell states [30,68–71]. The well-characterized trans-differentiation process known as the epithelial-to-mesenchymal transition [30,68] (EMT) facilitates the conversion of static epithelial cells into migratory mesenchymal cells, while the reverse pathway, the mesenchymal-to-epithelial transition [72–76] (MET), reverts mesenchymal cells to an epithelial state. Both pathways play critical roles during tissue development, wound healing, and cancer development [30,68,69,77]. Intermediate states exhibiting features of both epithelial and mesenchymal cells have been documented [78–82], suggesting a continuum rather than discrete cell states along the epithelial-mesenchymal axis. To ascertain the position of ARPE-19 cells along this continuum, we explored the impact of expressing master regulator transcription factors associated with epithelial and mesenchymal states. We hypothesized that if ARPE-19 cells were genuinely mesenchymal, they would not readily transition to a further mesenchymal state through experimental manipulation but could be effectively induced into an epithelial state. To test this, we generated stable ARPE-19 and control MCF10A cell lines containing doxycycline (dox)-inducible versions of the EMT-inducing transcription factor Snail [83–86] or the MET-inducing transcription factor OVOL2 [75,87–90] (Fig. 3A), and monitored phenotypic alterations in each cell population following the ectopic expression of Snail or OVOL2. In all experiments, comparison of parental cells in the presence and absence of dox ruled out spurious effects due to doxycycline, which have been observed in some systems [91–93]. Parental ARPE-19 cells exhibited a spindle-like, mesenchymal morphology using phase-contrast microscopy, unaffected by treatment with 1 ug/ml dox for 4 days (Fig. 3B). Induction of Snail in ARPE-19 cells resulted in negligible change in cellular morphology, while the induction of OVOL2 led to the formation of clustered, cobblestone islands, a hallmark feature of differentiated epithelial cells (Fig. 3B). Conversely, parental or uninduced MCF10A cells displayed a cobblestone morphology, transitioning to a spindle-like morphology with prominent extensions following ectopic expression of Snail, indicative of their conversion to a mesenchymal state. Additionally, MCF10A cells transitioned to an exaggerated cobblestone-island morphology after induction of OVOL2 (Fig. 3B). These observations suggest that subconfluent ARPE-19 cells are closely aligned with a fully mesenchymal state on the epithelial-mesenchymal continuum, whereas MCF10A cells are situated closer to a fully epithelial state, with some capacity to transition to an exaggerated epithelial state.

**Figure 3.**
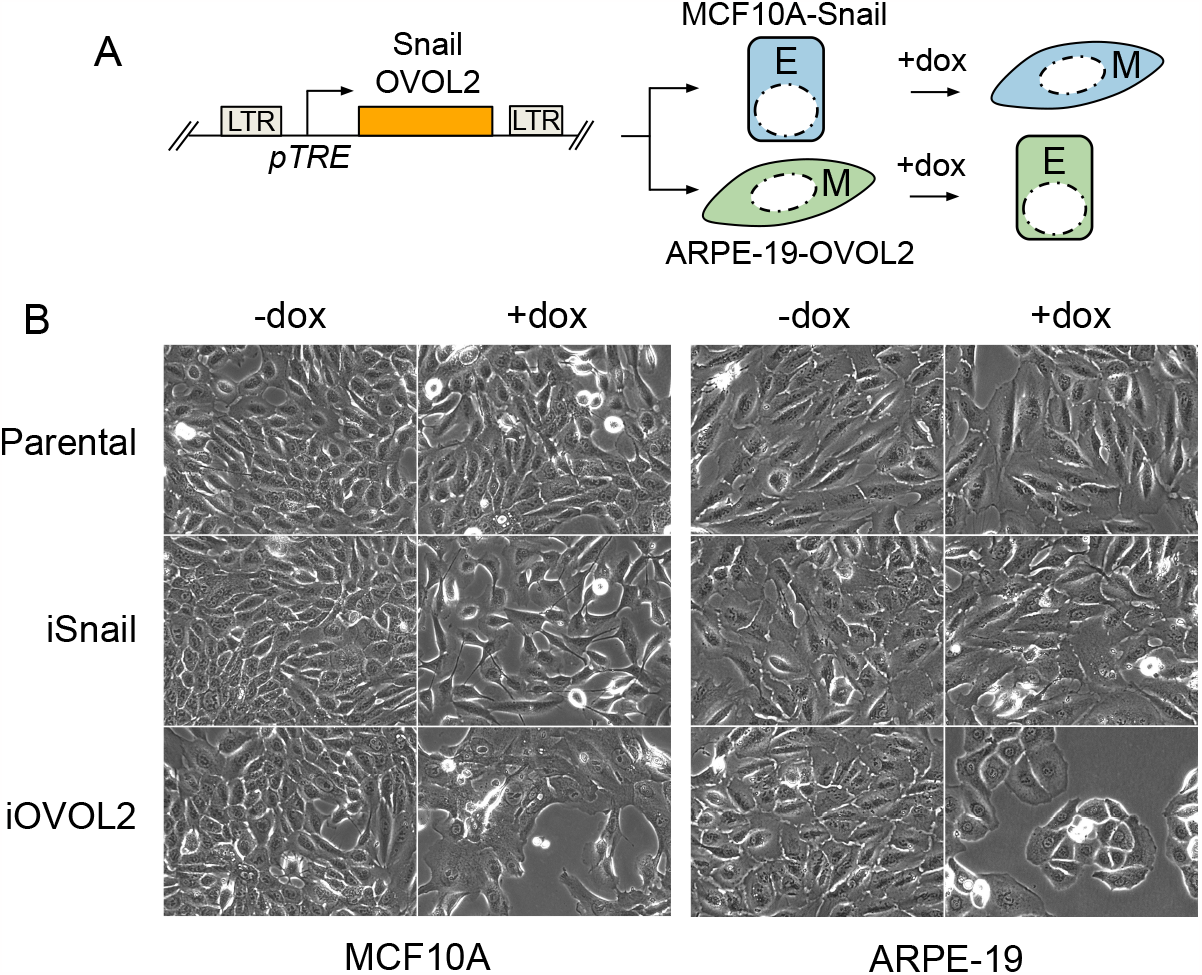
ARPE-19 Cells Show Morphological Features of Mesenchymal Cells and Can Be Driven to an Epithelial State Upon Expression of OVOL2. (A) Genetic system for assessing the impact of conditionally expressing EMT-inducing and MET-inducing transcription factors. The transcription factors Snail or OVOL2 are cloned into a doxycycline-inducible (dox), all-in-one lentiviral vector. Stable cell lines are established in MCF10A (MCF10A-i*GeneX*) or ARPE-19 (ARPE19-i*GeneX*) cells. Treatment with 1 ug/ml dox for 2 - 4 days drives cells toward a mesenchymal (Snail) or epithelial (OVOL2) cell fate and effects are observed. (B) Phase contrast images of showing morphological changes observed after 4 days of dox (1 ug/ml) treatment. MCF10A cells transition from a cobblestone epithelial morphology to a spindly mesenchymal morphology after expression of Snail. Parental ARPE19 cells display a mesenchymal morphology and show little-to-no morphological change after conditional expression of Snail, but can be driven towards a cobblestone epithelial morphology upon ectopic expression of OVOL2.

To substantiate these findings, we assessed changes in epithelial and mesenchymal gene expression patterns in dox-treated (dox+) and vehicle-treated (dox-) cells using *q*RT-PCR (Fig. 4A). MCF10A-iSnail cells treated with dox exhibited markedly reduced expression (16 to 32-fold) of epithelial genes (*CDH1, EpCAM, GJB3, MARVELD3, ST14*) and increased expression (up to 64-fold) of mesenchymal genes (*VIM, FN1, FBN1*) (Fig. 4A). Conversely, MCF10A-iOVOL2 cells treated with dox displayed a modest elevation in epithelial gene expression (4 to 16-fold) and a decrease in mesenchymal gene expression (4 to 9-fold) (Fig. 4B). ARPE-19-iSnail cells exhibited no statistically significant alterations in mesenchymal or epithelial gene expression upon dox treatment (Fig. 4C). However, ARPE-19-iOVOL2 cells treated with dox showed a substantial increase in epithelial gene expression (e.g., 64-fold for *CDH1* or 128-fold for *EPCAM*) and reduced expression of mesenchymal genes such as *VIM* and *FBN1* (Fig. 4D). Western analysis of E-cadherin, EpCAM, and Vimentin corroborated that changes in epithelial and mesenchymal gene expression were mirrored at the protein level (Figs. 4E and F). These data confirm that ARPE-19 cells, when cultured under sub-confluent conditions, exhibit a mesenchymal phenotype with a high capacity to trans-differentiate into an epithelial state, contrary to MCF10A cells, which display a strong epithelial phenotype with potent trans-differentiation capacity towards a mesenchymal state. Combined with our computational analyses, these results affirm that ARPE-19 cells cultured under low-confluency conditions possess mesenchymal features, as opposed to epithelial features.

**Figure 4.**
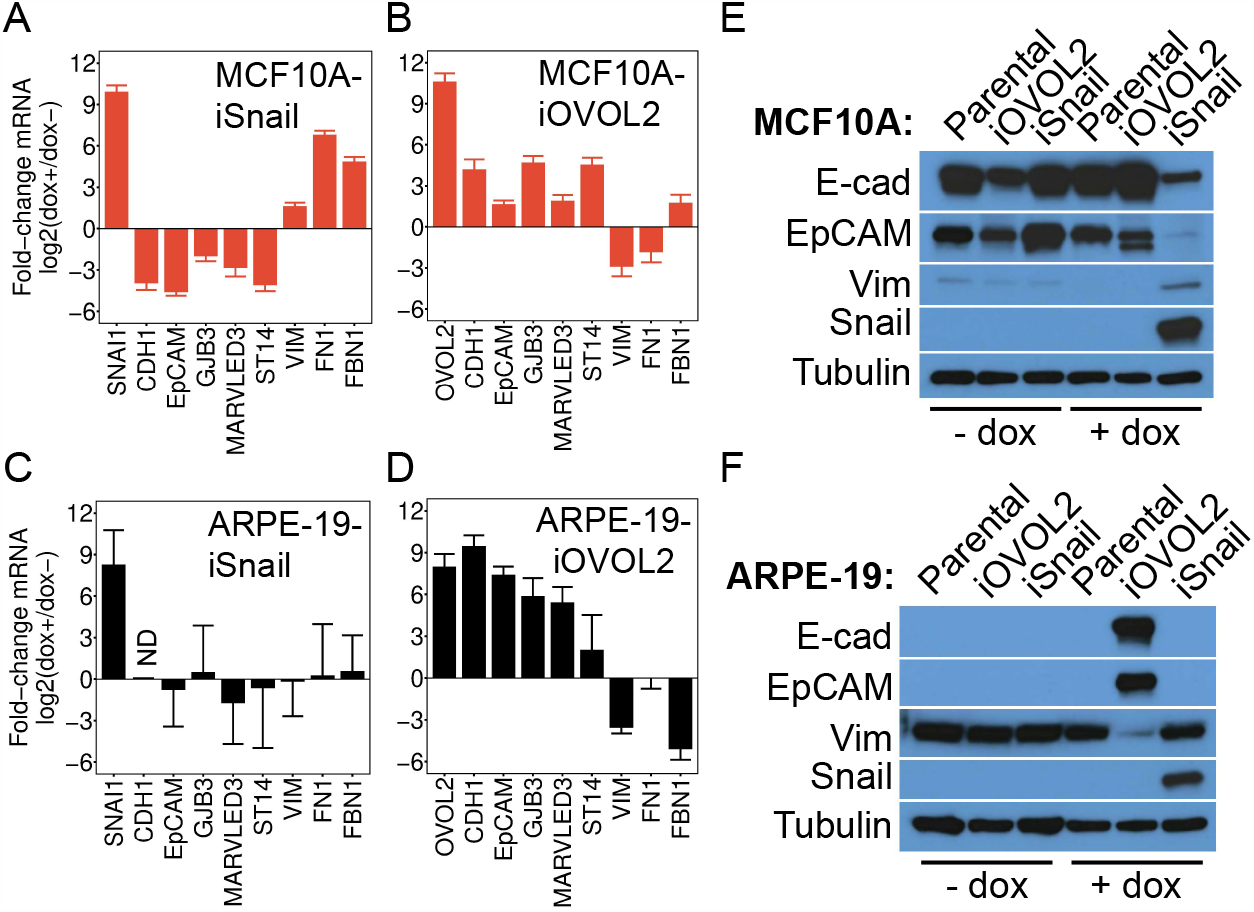
ARPE-19 Cells Can Be Experimentally Driven to an Epithelial Cell State by OVOL2, but Cannot Be Driven to a Further Mesenchymal State by Snail. Relative mRNA levels of epithelial (*CDH1, EpCAM, GJB3, MARVELD3, ST14)* and mesenchymal (*VIM, FN1, FBN1*) marker genes in MCF10A-iSnail (A), MCF10A-iOVOL2 (B), ARPE-19-iSnail (C), and ARPE-19-iOVOL2 (D) cells. Data were acquired using qRT-PCR and represent the fold-change in transcript levels after 4 days of treatment with 1 ug/ml dox or vehicle (water). Data represent the mean +/-SD from three biological replicates assayed in technical triplicate. *CDH1* was undetectable (“*ND”* = not determined) in the experiment in panel C, before or after treatment with dox. (E and F) Western analysis of epithelial (E-cad, EpCAM) and mesenchymal (VIM) proteins in parental, iSnail, and iOVOL2 MCF10A (E) or ARPE-19 cells (F). β-Tubulin was used as a loading control.

### Extended Confluent Culture Induces Epithelial and RPE-Specific Gene Expression in ARPE-19 Cells

Several studies have suggested that subculturing primary RPE [94,95] or ARPE-19 cells [23,95] at low confluency encourages an epithelial-to-mesenchymal transition (EMT). These studies demonstrated that the mesenchymal state associated with serial passaging could be reversed through extended maintenance (≥ 3 weeks) at high confluency [23] or transcription factor reprogramming [95]. This process caused increased expression of epithelial, visual-cycle, and RPE-signature genes [23,95]. We validated the induction of epithelial, visual-cycle, and RPE-signature genes in our lab stock of ARPE-19 cells over a three-week time course (Fig. 5A and B). Interestingly, some mesenchymal genes such as *FN1* and *FBN1* were also elevated during this period (Fig. 5A), suggesting that long-term cultured ARPE-19 cells may not fully transition to an epithelial state, but rather may attain a hybrid state embodying characteristics of both epithelial and mesenchymal cell states. Alternatively, even longer periods of confluent culture may be required to transition ARPE-19 cells to a fully differentiated epithelial cell state. Supporting this possibility, we were unable to detect the expression of the rate-limiting visual cycle enzyme RPE65, known to be induced 10,000-fold following four months of high-confluency culture [23].

**Figure 5.**
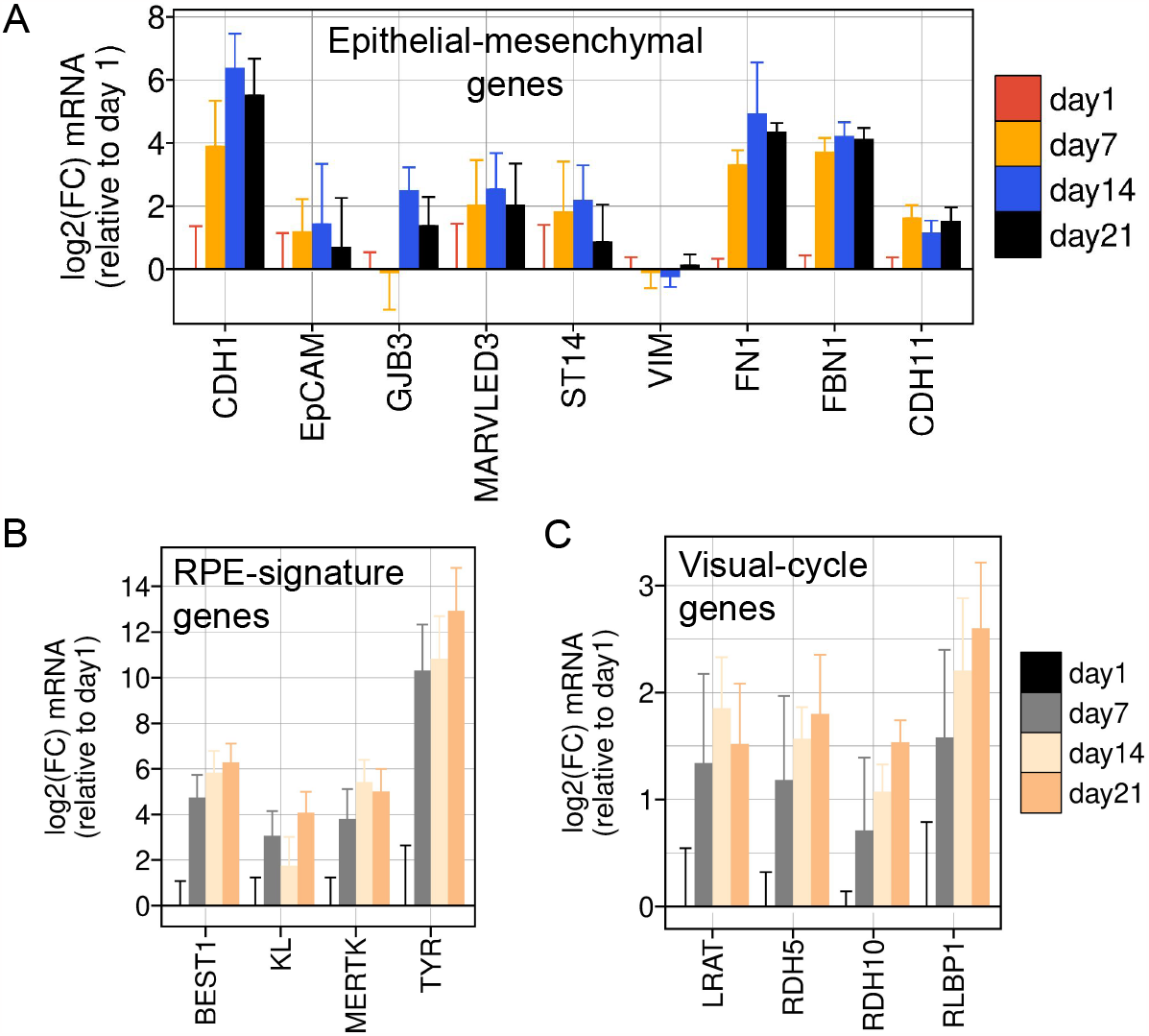
Gene Expression Changes Induced by Long-Term Culture of ARPE-19 Cells. Gene expression changes induced in ARPE-19 cells over three weeks of high confluency culture. Total RNA was isolated at the indicated time points, reverse-transcribed, and relative mRNA levels were measured using quantitative RT-PCR (*q*RT-PCR). (A) Relative mRNA levels of epithelial (*CDH1, EpCAM, GJB3, MARVELD3, ST14)* and mesenchymal (*VIM, FN1, FBN1*) genes. (B) Relative mRNA levels of RPE-signature genes. (C) Relative mRNA levels of visual cycle genes responsible for retinol metabolism. Data represent the mean +/-SD from three biological replicates assayed in technical triplicate.

**Figure 6.**
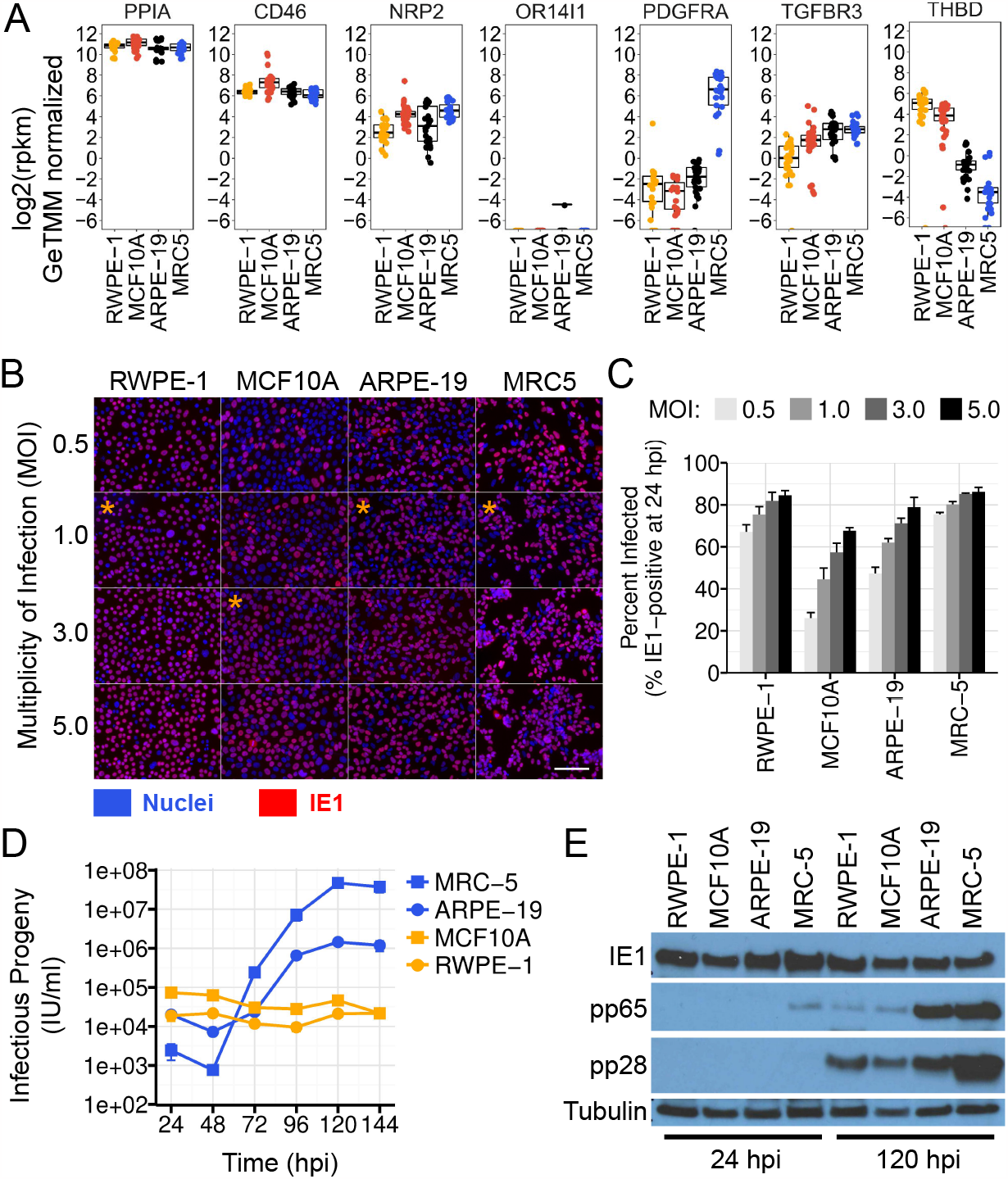
ARPE-19 Cells Phenocopy Fibroblasts in Experimental Infection Assays with Human Cytomegalovirus. (A) Absolute RNA-expression levels of HCMV entry factors across ARPE-19, MCR5, RWPE-1, and MCF10A cells. Data are normalized public gene expression data from Figures 1 and 2. PPIA RNA levels are shown as a normalization control. (B and C) HCMV susceptibility analysis. Fluorescence microscopy images of different cell types infected with HCMV TB40-BAC4 at various MOIs. IE1 (*red*) was visualized at 24 hpi by indirect immunofluorescence and nuclei were counterstained with DAPI (*blue*). Orange asterisks mark the MOIs used for growth curve analysis in panel *D. Scale bar;* 100 um. The percentage of infected cells (C) was quantified for each cell type and multiplicity using Cellprofiler software (see “Materials and Methods”). Data are mean +/-SD of three independent infections. (D) Single-step growth analysis of HCMV across four cell types. Infectious progeny released into the cell supernatants was assayed by IU-assay over a six day time course. MRC-5, ARPE-19, and RWPE-1 cells were infected at 1 IU/cell. MCF10A cells were infected at 3 IU/cell. Data are mean +/-SD of three independent infections per cell type, per time point. In most cases, error bars are smaller than the data point symbols. (E) Immunoblot of HCMV proteins at 24 and 120 hpi, in infected epithelial and mesenchymal cell lines. Tubulin is shown as a loading control.

### ARPE-19 Cells Phenocopy Fibroblasts in Experimental Infection Assays with Human Cytomegalovirus

Given our observations regarding the mesenchymal phenotype of ARPE-19 cells, we hypothesized that HCMV might behave differently in ARPE-19 than strongly epithelial cell lines in experimental infection studies. To test this possibility, we first determined the susceptibility of the epithelial cell lines MCF10A and RWPE-1 to HCMV infection. HCMV utilizes two glycoprotein complexes, termed “trimer” (gH, gL, gO) and “pentamer” (gH, gL, UL128, UL130, and UL131), to bind cellular entry receptors on various cell types [11,16,96–101]. ARPE-19, MCF10A, and RWPE-1 cells express low levels of PDGFRα mRNA (Fig. 6A), the cellular receptor for the HCMV trimer. However, they do express entry factors for pentamer, including NRP2, TGFBR3, and CD46 (Fig. 6A), suggesting that entry into these cell lines requires pentamer-containing strains of HCMV. Therefore, for this analysis, we employed HCMV strain TB40-BAC4 [35], which retains epithelial tropism and high levels of pentamer on virions when grown in ARPE-19 cells [10,11]. To assess susceptibility, we infected each cell line with HCMV at different MOIs and monitored IE1 expression at 24 hpi, a surrogate for viral entry. MRC-5 fibroblasts were used as fully susceptible control cells in these experiments. ARPE-19, MRC-5, and RWPE-1 cells were all highly susceptible to TB40-BAC4, with infection rates close to Poisson-theoretical (Figs. 6B and C). MCF10A cells showed slightly reduced susceptibility, but greater than 50% of the cells could be infected at a multiplicity of 3 IU/cell (Figs. 6B and C).

To assess permissivity and infectious progeny production, we next infected each cell line and monitored viral protein biosynthesis and infectious progeny production over a six day infection time course. Each cell population was infected with an identical stock of HCMV TB40-BAC4 at a multiplicity of 1 IU/cell, with the exception of MCF10A cells, which were infected at 3 IU/cell to compensate for their reduced susceptibility. ARPE-19 cells produced logarithmically increasing yields of cell-free infectious progeny with similar kinetics to fully productive MRC-5 fibroblasts, although peak progeny yields were diminished by approximately 50-fold (Fig. 6D). In contrast, MCF10A and RWPE-1 cells showed a reproductive index near unity, with only residual input infectivity recovered from the cell supernatants after six days of monitoring (Fig 6D). Analysis of steady-state viral protein levels in the infected cells from each population showed that the immediate-early protein IE1 was expressed at similar levels in all cell lines at 24 hpi, but late protein production was diminished late during infection (120 hpi), specifically in the epithelial cell lines (Fig. 6D). Steady-state levels of the true-late protein pp28 were modestly reduced in RWPE-1 and MCF10A cells, while expression of the leaky-late protein pp65 was dramatically reduced (Fig 6D). It is not clear whether these alterations result from decreased mRNA or protein synthesis, or decreased stability. However, their occurrence specifically in the cell lines with strong epithelial character, which fail to produce infectious progeny, would seem to indicate the presence of cell-type specific mechanisms that strongly influence viral biosynthesis and/or spread.

Overall, these data show that HCMV behaves quite differently in ARPE-19 cells than strongly epithelial cell lines like MCF10A and RWPE-1. They also suggest that HCMV may establish a non-canonical pattern of infection in some epithelial cell types, where biosynthesis is atypical, or delayed; or, where the virus is transmitted without the production of cell-free progeny.

## Discussion

The behavior of HCMV in epithelial cells has been a topic of interest in the HCMV field for some time. Early studies of HCMV infection in ARPE-19 or telomerase-immortalized RPE cells observed exclusive cell-associated replication, with little-to-no release of cell-free progeny and a low reproductive index (the ratio between input and output virus) [102,103]. However, these studies used HCMV propagated in human fibroblasts, which is known to decrease epithelial tropism [10,11,104–106]. Growth of clinical HCMV strains in ARPE-19 cells yields virions with epithelial-tropic properties, such as retention of high levels of the pentamer glycoproteins on virions [11] and a shift in entry route from endocytosis to direct-cell fusion [18]. These changes facilitate entry into a variety of epithelial and myeloid cell-types [10,11], as well as the production of high yields of cell-free progeny in ARPE-19 cells [10]. A similar effect on HCMV tropism has been observed for stocks propagated in endothelial cells [35,106]. By preparing epithelial-tropic stocks of HCMV in ARPE-19 cells, we were able to compare HCMV permissivity across epithelial and mesenchymal cell lines using an identical virus preparation. We observed a strikingly different pattern of HCMV infection in mesenchymal and epithelial cells, with ARPE-19 cells behaving like productive fibroblasts, and epithelial cells displaying an atypical, potentially abortive, infection. The consequences and generality of this phenomenon are currently unknown. However, our realization that low confluency ARPE-19 cultures typically lose their epithelial characteristics redirects future studies towards alternative epithelial cell models.

To the best of our knowledge, this is the first study to utilize MCF10A or RWPE-1 cells for experimental infection with HCMV. Both mammary epithelial cells and prostate epithelial cells are physiological sites of HCMV infection [7,61,62]. Therefore, these cell lines could be useful in future studies aimed at understanding HCMV replication in these cell types, both of which may harbor persistent HCMV *in-vivo*.

ARPE-19 cells are a commonly used epithelial infection model for HCMV [12,18,107–122], as well as other human herpesviruses (e.g. VZV or HSV-1) [123–125], filoviruses (e.g. Ebola) [126], flaviviruses (e.g. Zika or Dengue) [127–129], and coronaviruses (e.g. SARS-CoV-2) [130]. Despite a number of studies showing that ARPE-19 requires prolonged periods of contact inhibited cell culture for functional differentiation [9,23], the use of undifferentiated, low-confluency ARPE-19 cultures (95% to just confluent) for infection studies is common. For example, we could find only one study in the HCMV literature explicitly stating that long-term, high confluency cultures were utilized [22]; a study specifically designed to assess the effects of epithelial polarization on HCMV infection. The capacity of ARPE-19 to de-differentiate to a mesenchymal cell state in a density dependent manner remains an underappreciated aspect of this cell line.

The data in this study provide clear evidence that low confluency ARPE-19 cultures establish a mesenchymal, rather than epithelial, cell state. Using public gene expression data, we also provide evidence that this phenomenon pervades across a variety of studies using ARPE-19, suggesting that the loss of epithelial characteristics at low confluency is a general feature of the cell line, rather than a specific property of cultures maintained in our laboratory. Hopefully this work will increase awareness of the cell state plasticity inherent to the ARPE-19 cell line, which continues to be an important *in-vitro* model for studying retinal biology. Is HCMV infection sensitive to the epithelial-mesenchymal cell state of cells? The answer to this question remains unclear. However, combining our observation that HCMV progeny production is restricted in some epithelial cell-types with our previous data showing that HCMV infection itself can induce epithelial gene expression [10], it is possible that one or more feature of the epithelial cell state program is intrinsically restrictive to HCMV, or alters the mode of transmission from cell-free to cell-associated. Alternatively, epithelial-specific transcriptional or translational mechanisms could control the kinetics of HCMV biosynthesis, potentially benefiting the virus by coordinating biosynthesis with intrinsic immune detection. Such a mechanism might explain the slow, persistent, infections thought to occur in some epithelial tissues. Modulating epithelial cell state features via the epithelial-to-mesenchymal transition (EMT) and observing the effects on HCMV biosynthesis and spread could be useful for studying how epithelial-mesenchymal cell states impact the HCMV infectious cycle. Ultimately, however, more sophisticated models will be required to determine whether epithelial-mesenchymal cell state dynamics influence natural infections *in-vivo*.

One limitation of our study is that we have only assessed HCMV infection in a small number of epithelial cell lines. It will be important to extend these observations to additional epithelial cell types in the future, to determine if our results generalize across diverse epithelial cell types or are specific to the particular mammary and prostate cell lines we have tested.

In summary, our results indicate that subconfluent ARPE-19 cells are not ideal for studying HCMV interactions with epithelial cells *in-vitro*, and identify the epithelial-mesenchymal cell state axis as a potential regulator of HCMV infection in epithelial cells. Our findings also underscore the necessity for caution when utilizing density-dependent cell lines such as ARPE-19 for cell biological experiments. We hope that our work serves to inform future studies utilizing ARPE-19 cells as a model for epithelial infections, not only with HCMV, but also with other human pathogens.

## Acknowledgments

We thank Dr. Alan McLachlan, PhD (UIC) for critical reading of our manuscript and general support, Dr. Thomas Shenk, PhD (Princeton University) for advice and providing the HCMV monoclonal antibodies used in this study, and the UIC College of Medicine for providing funding for this research.

## Supplementary Data

Table S1 - Public RNA-seq metadata used in this study.

Table S2 - *q*RT-PCR Primers used in this study.

Table S3 - Antibodies used in this study.

