## Supplementary material for "Subconfluent ARPE-19 Cells Display Mesenchymal Cell-State Characteristics and Behave Like Fibroblasts, Rather than Epithelial Cells, in Experimental HCMV Infection Studies": Table S2

**Table S2.** ***q*****RT-PCR primers used in this study.**

| Primer name | Sequence | Source |
|---|---|---|
| VIM-fwd | GAGAACTTTGCCGTTGAAGC | (Mani et al., 2008) |
| VIM-rev | GCTTCCTGTAGGTGGCAATC | (Mani et al., 2008) |
| CDH1-fwd | TGTTCACCATTAACAGGAACAC | This study |
| CDH1-rev | GGGTATACGTAGGGAAACTCTC | This study |
| CDH2-fwd | TCAGTGAAGGAGTCAGCAG | This study |
| CDH2-rev | CTTCTGCCTTTGTAGGTGG | This study |
| CDH11-fwd | GCTGACTTGTGAATGGGAC | This study |
| CDH11-rev | TTGAGCTCATCACGTCAGG | This study |
| FN1-fwd | AAATGGCCAGATGATGAGC | This study |
| FN1-rev | TAACACGTTGCCTCATGAG | This study |
| FBN1-fwd | TAGGATGTGCAAAGATGAGG | This study |
| FBN1-rev | ATGAGGTTCTTGCATTCCA | This study |
| PPIA-fwd | AGCCAGGTACTTGGTGCTACAGTC | This study |
| PPIA-rev | TGCAGGTAGTCTGCGCCTTAAC | This study |
| C1orf116-fwd | CTCTGTCTCCATCTCTGCC | This study |
| C1orf116-rev | GCTATCACTCTCCACTGGG | This study |
| EPCAM-fwd | CGAGTGAGAACCTACTGGA | This study |
| EPCAM-rev | TGATCTCCTTCTGAAGTGCA | This study |
| GJB3-fwd | CTCATCATTGAGTTCCTCTTCC | This study |
| GJB3-rev | GCATATTGAAGCCATGCCA | This study |
| MARVELD3-fwd | AGAGATATCTGCCCTCGAC | This study |
| MARVELD3-rev | TCTGACTGGTAATATTCCACCTC | This study |
| ST14-fwd | CATGGAACATTGAGGTGCC | This study |
| ST14-rev | ATCTCCACGTAGTCCTTGG | This study |
| OVOL2-fwd | AAATCAAGTTCACCACAGGC | This study |
| OVOL2-rev | ACTTGAGGTGACGGTTCAG | This study |
| SNAI-fwd | TCTTTCCTCGTCAGGAAGC | This study |
| SNAI-rev | AGGTAAACTCTGGATTAGAGTCC | This study |
