## Supplementary material for "Subconfluent ARPE-19 Cells Display Mesenchymal Cell-State Characteristics and Behave Like Fibroblasts, Rather than Epithelial Cells, in Experimental HCMV Infection Studies": Table S3

**Table S3. Antibodies used in this study.**

| Antibody | Dilution | Source | Identifier |
|---|---|---|---|
| Mouse monoclonal anti-E-Cadherin | 1:500 | BD Biosciences | Cat# 610181; RRID: AB_397580 |
| Goat polyclonal anti-EPCAM | 1:500 | R & D Systems | Cat# AF960; RRID: AB_355745 |
| Mouse monoclonal anti-β-tubulin | 1:2000 | The Developmental Studies Hybridoma Bank | Cat# E7; RRID: AB_528499 |
| Mouse monoclonal anti-Vimentin | 1:1000 | The Developmental Studies Hybridoma Bank | Cat# AMF-17b; RRID: AB_528505 |
| Goat polyclonal anti-Snail | 1:500 | R & D Systems | Cat# AF3639; RRID: AB_2191738 |
| Rabbit polyclonal anti-OB-Cadherin | 1:500 | Cell Signaling Technology | Cat# 4442; RRID: AB_10547881 |
| Mouse monoclonal anti-N-Cadherin | 1:500 | Cell Signaling Technology | Cat# 14215; RRID: AB_2798427 |
| Mouse monoclonal anti-IE1 (clone 1B12) | 1:1000 | Shenk lab, Princeton University (Zhu et al., 1995) | N/A |
| Mouse monoclonal anti-UL99/pp28 (clone 10B4-29) | 1:1000 | Shenk lab, Princeton University (Silva et al., 2003) | N/A |
| Mouse monoclonal anti-UL83/pp65 (clone 8F5) | 1:1000 | Shenk lab, Princeton University (Nowak et al., 1984) | N/A |
